# Proximity labeling reveals differential interaction partners of the human mitochondrial import receptor proteins TOMM20 and TOMM70

**DOI:** 10.1101/2024.10.25.620316

**Authors:** Saira Akram, Katharina I. Zittlau, Boris Maček, Ralf-Peter Jansen

**Author notes:** Corresponding author: Interfaculty Institute of Biochemistry, University of Tübingen, Auf der Morgenstelle 34, 72076 Tübingen, Germany.

## Abstract

Import of most mitochondrial proteins requires that their precursor proteins are bound by the (peripheral) receptor proteins TOM20, TOM22, and TOM70. For budding yeast TOM20 and TOM70, there is evidence of specific yet overlapping substrate recognition, but no such data is available for metazoan cells. Using APEX2-based proximity labeling, we thus created association profiles for human TOMM20 and TOMM70 in HeLa cells. We particularly focused on their interaction with RNA-binding proteins (RBPs) since there is evidence for RNA association with the mitochondrial outer membrane (MOM) and local translation at the mitochondrial surface, but these processes are poorly understood. Our results show a preferred association of several RBPs and translation factors with TOMM20 over TOMM70. These include SYNJBP2, a previously identified membrane-bound RBP that binds and protects mRNAs encoding mitochondrial proteins. Translational inhibition by puromycin resulted in an even increased association of these RBPs with TOMM20 compared to TOMM70, suggesting that TOMM20 but not TOMM70 might play a role in preserving cellular hemostasis during translation stress by retaining protective RBPs and translation-related proteins at the MOM.

## Introduction

Protein sorting to intracellular organelles is an essential process in eukaryotic cells since the majority of proteins are nuclear-encoded but play physiological roles in specific cellular compartments. Mitochondria must import more than 1000 different preproteins and sort them to the various intramitochondrial compartments. These preproteins contain specific targeting signals that are recognized by the mitochondrial import machinery ^1^. A major entry gate is the translocase of the outer mitochondrial membrane (TOM). This protein complex contains the translocation pores formed by the TOM40 (TOMM40 in mammals) subunit and a central preprotein receptor subunit, TOM22/TOMM22. TOMM22 accepts its substrates from two additional receptor subunits, TOM20/TOMM20 and TOM70/TOMM70, and transfers them to the translocating TOMM40 pore. TOMM20 and TOMM70 recognize the targeting signals but differ in their substrate specificity, although overlap can be observed^2,3^. Whereas TOMM20 binds to preproteins targeted to the inner mitochondrial membrane (MIM) and the matrix, TOMM70 prefers substrates that are alpha-helical proteins destined to the outer or inner mitochondrial membrane^4–6^.

Though both receptors are dynamically associating with TOMM22/TOMM40 subcomplex, it is considered that TOMM70 is more loosely associated with the core complex compared to TOMM20 ^7–9^. TOMM70 typically migrates as a homodimer on Blue Native PAGE, indicating its less stable interaction with the core complex^10,11^. Conversely, TOMM20, when solubilized with mild detergents, also exhibits a loose association with the core complex but migrates as a high-molecular-weight complex above 400 kDa in Blue Native PAGE, distinct from TOMM70^12–14^. Delivery of substrate proteins to the receptors involves chaperones that prevent translated precursor proteins from aggregation and can target them to the corresponding receptors^15^.

Consistent with the proposed role of chaperones, *in vitro* import assays suggest that mitochondrial protein import occurs post-translationally. However, increasing evidence also suggests local translation of mRNAs at the mitochondrial surface and co-translational import of mitochondrial proteins^16–18^. This evidence includes the presence of ribosomes at the mitochondrial outer membrane (MOM)^19^, co-fractionation of thousands of mRNAs with yeast mitochondria ^20^, or the identification of mRNAs by Ribo-Seq from ribosomes that were labeled by a biotin ligase targeted to the MOM^21^. Translation-dependent accumulation of mRNAs at the MOM has also been reported from mammalian HEK293 cells^16^. Here, proximity-labeling based RNA sequencing identified hundreds of mRNAs at the MOM, with the majority being mRNAs encoding mitochondrial proteins (mitoRNAs). Treatment with the translation inhibitor puromycin causes detachment of most of these mRNAs from ribosomes, suggesting that their localization depends on translation, most likely via the Mitochondrial Targeting Signal (MTS) in the nascent chain complex^22,23^. Translation of mRNAs might occur locally at the TOM complex with some of its subunits participating in the co-translational import as previous studies have shown that yeast TOM20 contributes to the co-translational import of the mitochondrial proteins^18^. In parallel, TOM70 may also contribute to some level to localized translation at the MOM, as TOM70 depletion in yeast and mammalian cells led to a reduction in the levels of mitochondrial-localized mRNAs^18,24^, or to the dissociation of ribosomes associated with a subset of mitoRNAs^17^.

Furthermore, RNA-binding proteins (RBPs) have been identified in yeast, *Drosophila*, and mammalian cells that not only bind to mRNAs encoding mitochondrial proteins but control stability, translation, or localization of these mRNAs^22^. In budding yeast, the loss of RBP Puf3p results in de-localization of mRNAs to the MOM^18,25^. The protein is not only involved in mRNA localization to the MOM but might also block translation of these mRNAs while in transit to the MOM. In mammals, the RBPs Clustered mitochondria homolog (CLUH) and Synaptojanin 2 binding protein (SYNJ2BP) have been extensively studied. CLUH preferentially binds mRNAs of nuclear-encoded mitochondrial proteins and its depletion results in reduced translation of proteins encoded by these mRNAs, resulting in defects in mitochondrial morphology^26^. SYNJBP2 has been identified as a component of RNA-protein complexes at the MOM and is essential for localization of its target mRNAs^27^ and piggy-back traveling of *PINK1* mRNA with mitochondria in neurons, while SYNJ2BP knockdown redistributes *PINK1* mRNA into RNA granules and inhibits local mitophagy^28^. It specifically anchors its target mRNAs at the MOM under translation stress, facilitating their local translation and further import into mitochondria. Furthermore, its loss in HEK293 cells compromises the function of the OXPHOS (oxidative phosphorylation) complex^27^.

To compare the local interactome of TOMM20 and TOMM70, and with a specific focus on RBPs associated with the two receptors, we applied an APEX2 based proximity labeling approach and fused the APEX2 enzyme to either TOMM20 or TOMM70 in HeLa cells. We identified distinct sets of associated proteins revealing that each receptor subunit interacts with its own unique set of proteins. Among these proteins are translation factors and ribosomal proteins but also RBPs like SYNJ2BP that specifically associate with TOMM20, which indicates an association of the translation machinery with this TOM receptor subunit and suggests that TOMM20 is more engaged than TOMM70 in localized translation at MOM.

## Results

### Expression of functional TOMM20-APEX2 and TOMM70-APEX2 fusion proteins

To characterize the interactomes of the two main receptor subunits of the human TOM complex, we generated expression constructs targeted for integration into the HeLa-EM2-11ht genome (see Methods). In both hybrid proteins, the APEX2 proximity labeling enzyme^29^ is located at the cytoplasmic carboxyl (C-) terminus of the respective TOMM protein, separated by a short linker sequence (GGSGDPPVAT). Both fusion proteins contain an additional V5 epitope tag^30^ for detection (**Figure 1A**) and are expressed from an inducible pTet promoter by addition of doxycycline to the medium. We initially used the V5 tag to test if the fusion proteins are targeted to mitochondria (**Figure 1B**). After 24 h induction, immunofluorescence microscopy at super resolution revealed a co-localization of TOMM20-APEX2-V5 or TOMM70-APEX2-V5 with a mitochondrial marker (TOMM22), which indicates the correct targeting of the fusion proteins. Furthermore, mitochondrial morphology remains unchanged, which suggests that the fusion proteins have no negative impact on mitochondrial function. To test for correct association of TOMM20-APEX2 with the TOM complex we performed pull down assays with endogenous TOM complex subunits. TOMM20-APEX2 expression was induced for 24 hours prior to lysis, mitochondria were isolated and solubilized with a digitonin-containing buffer. Detergent extracts were subjected to co-immunoprecipitation using an anti-TOMM22 or anti-TOMM40 antibody and the fusion proteins detected via the V5 tag. The 49 kDa TOMM20-APEX2 protein fusion was found in both immunoprecipitates (**Fig 1C and 1D**). In a reciprocal co-immunoprecipitation experiment, using anti-V5 beads we co-purified TOMM22, as shown by Western blotting using an anti-TOMM22 antibody (**Fig 1E**). These experiments confirmed that TOMM20-APEX2 associates with both members of the endogenous TOM complex. We also analyzed integration of both fusion proteins into mitochondria by subcellular fractionation. Like endogenous TOMM40 and TOMM20, TOMM20-APEX2 (detected by an anti-TOMM20 antibody) is highly enriched in the mitochondrial fraction, whereas glyceraldehyde-3-phosphate dehydrogenase (GAPDH) is primarily detected in the cytoplasmic fraction **(Figure 1F)**. Subcellular co-fractionation was also used to assess distribution of TOMM70-APEX2. Detection of the fusion protein and the endogenous TOMM70 via an anti-TOMM70 antibody reveals similar distribution patterns in mitochondrial versus cytosolic fractions, indicative of correct targeting of the fusion protein to mitochondria (**Figure 1G**). Importantly, these experiments also revealed similar expression levels of endogenous TOMM20 or TOMM70 and the corresponding fusion proteins TOMM20-APEX or TOMM70-APEX, respectively.

**Figure 1.**
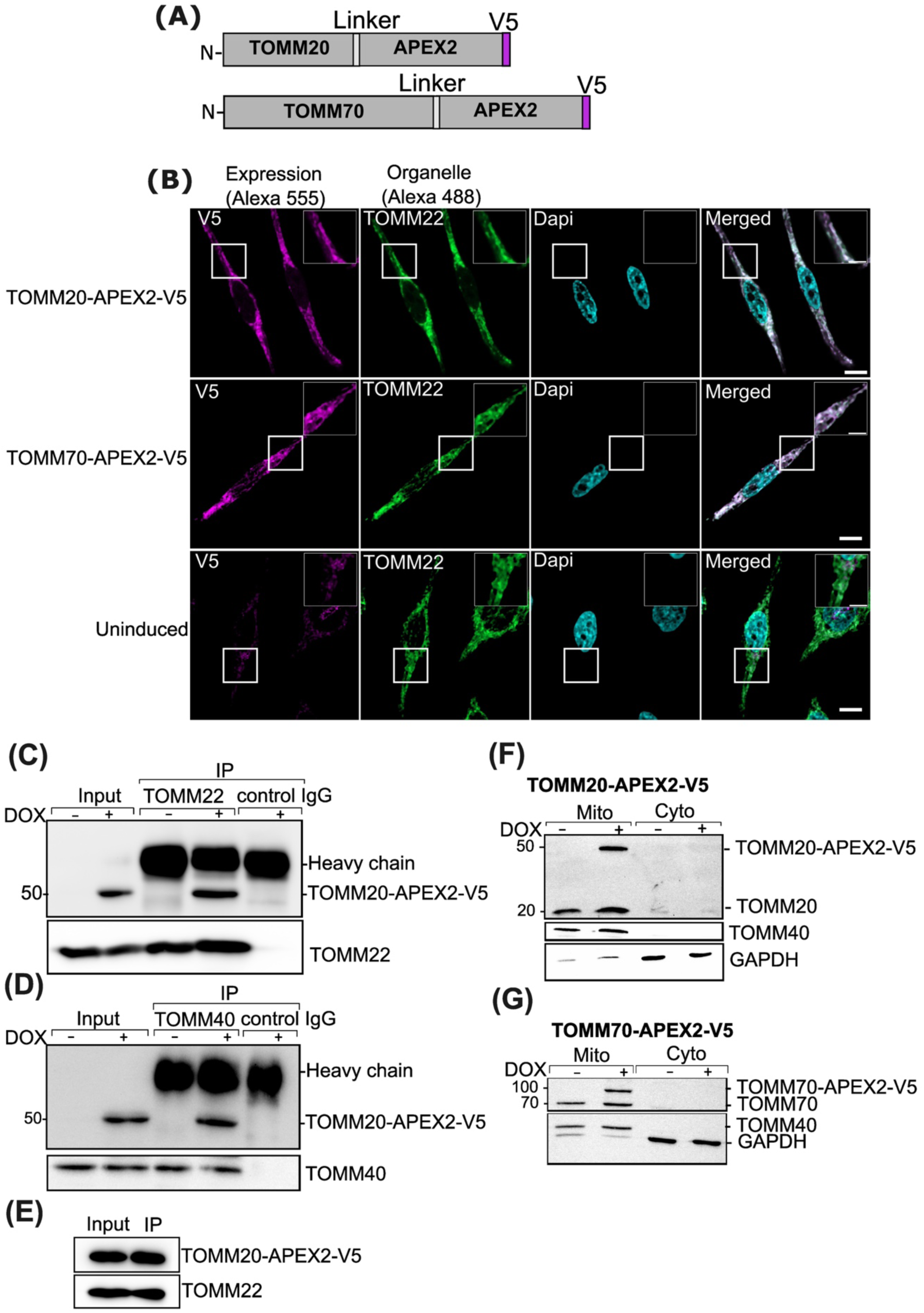
Functional expression of TOMM20– and TOMM70-APEX2 fusion proteins. (**A**) Constructs map of TOMM20-APEX2 and TOMM70-APEX2 fusion proteins. APEX2 is fused to the gene of the target protein at the C terminus via a short intervening linker sequence of 10-amino acids. **(B)** Confocal fluorescence imaging confirming the APEX2 localization in the cells stably expressing the indicated APEX2 fusion proteins. After 24 h induction with DOX and subsequent fixation, the fusion proteins are immunolabeled with antibody directed against V5 (Magenta). Endogenous TOM complex is visualized using anti-TOM22 antibody (Green) (Scale bar: 10 μm). Nuclei are stained with DAPI (cyan). Incents are the magnified portion of the cell showing on the uppermost right of each panel (Scale bar, 5 μm) **(C, D)** Western blot to show interaction of Tomm20-APEX2 fusion protein with endogenous TOM complex subunits analyzed by pull-down assay. Hela cells are induced with DOX for 24 hours to stably express TOMM20-APEX2 fusion protein prior to lysis. Mitochondria are isolated, solubilized with digitonin-containing buffer and further subjected to immunoprecipitation wherein the TOM complex is specifically pulled down either using TOMM22 IgG **(C)**, or TOMM40 IgG **(D)**. Eluates from IPs of uninduced cells (-) are also loaded in parallel. **(E)** Western blot showing the pulldown analysis of TOMM20-APEX2 fusion protein in isolated mitochondria using V5 beads. **(F)** Western blot analysis of mitochondrial and cytosolic fractions of TOMM20-APEX2 expressing cells. Cells are either not or induced with DOX for 24 hours, mitochondria are isolated and further solubilized with digitonin-containing buffer and isolated fractions are analysed with TOMM20 and TOMM40 antibody. GAPDH is used as cytosolic marker. Endogenous (∼20 kDa) and APEX2 containing TOMM20 (∼50 kDa) subunits are confirmed by TOMM20 antibody. **(G)** Western blot analysis of mitochondrial and cytosolic fractions of TOMM70-APEX2 expressing cells. Endogenous (70 kDA) and APEX2 containing TOMM70 (∼100 kDa) subunits are detected by TOMM70 antibody. Cytosolic fraction is validated by GAPDH.

### Specific biotinylation activity of TOMM20-APEX2 and TOMM70-APEX2

Since correct localization of the TOMM20– and TOMM70-APEX2 fusion proteins to mitochondria (**Figure 1**) as well as their co-purification with TOMM22 and TOMM40 proteins suggest that both APEX2 fusions are correctly integrated into the TOM complex, we next tested for biotinylation activity of the fusion proteins. Besides endogenous biotinylated proteins that can be detected under all conditions, additional biotinylation in cells with integrated TOMM20– or TOMM70-APEX2 constructs could only be detected upon expression of the respective protein (**Figure 2**, ‘DOX’), addition of biotin-phenol (‘BP’), and of H₂O₂(‘H₂O₂’). We also verified biotinylation by *in situ* labeling of biotinylated proteins, using neutravidin-Alexa647 (**Figure 3**). Even under conditions that do not allow APEX activity (e.g. no expression, no BP, no H₂O₂), a neutravidin-Alexa647 signal is detectable that overlaps with the mitochondrial location of the APEX2 fusion proteins. This most likely reflects detection of endogenous biotinylated proteins such as mitochondrial pyruvate carboxylase or acetyl-CoA-caboxylase^31^. Only HeLa cells expressing the TOMM20– and TOMM70-APEX2 fusion proteins show an additional, and stronger biotin-dependent fluorescence (**Figure 3A, B**, left column), indicating an active APEX2 enzyme. Surprisingly, the biotinylation seen in TOMM20-APEX and TOMM70-APEX expressing cells was not restricted to the mitochondrial location. However, this has been seen before for TOMM20-APEX2^32^ and has been explained to depend on the cytosolic orientation of the APEX2 part of the fusion protein, diffusion of the activated biotin-phenoxyl radical and labeling of cytosolic components.

**Figure 2.**
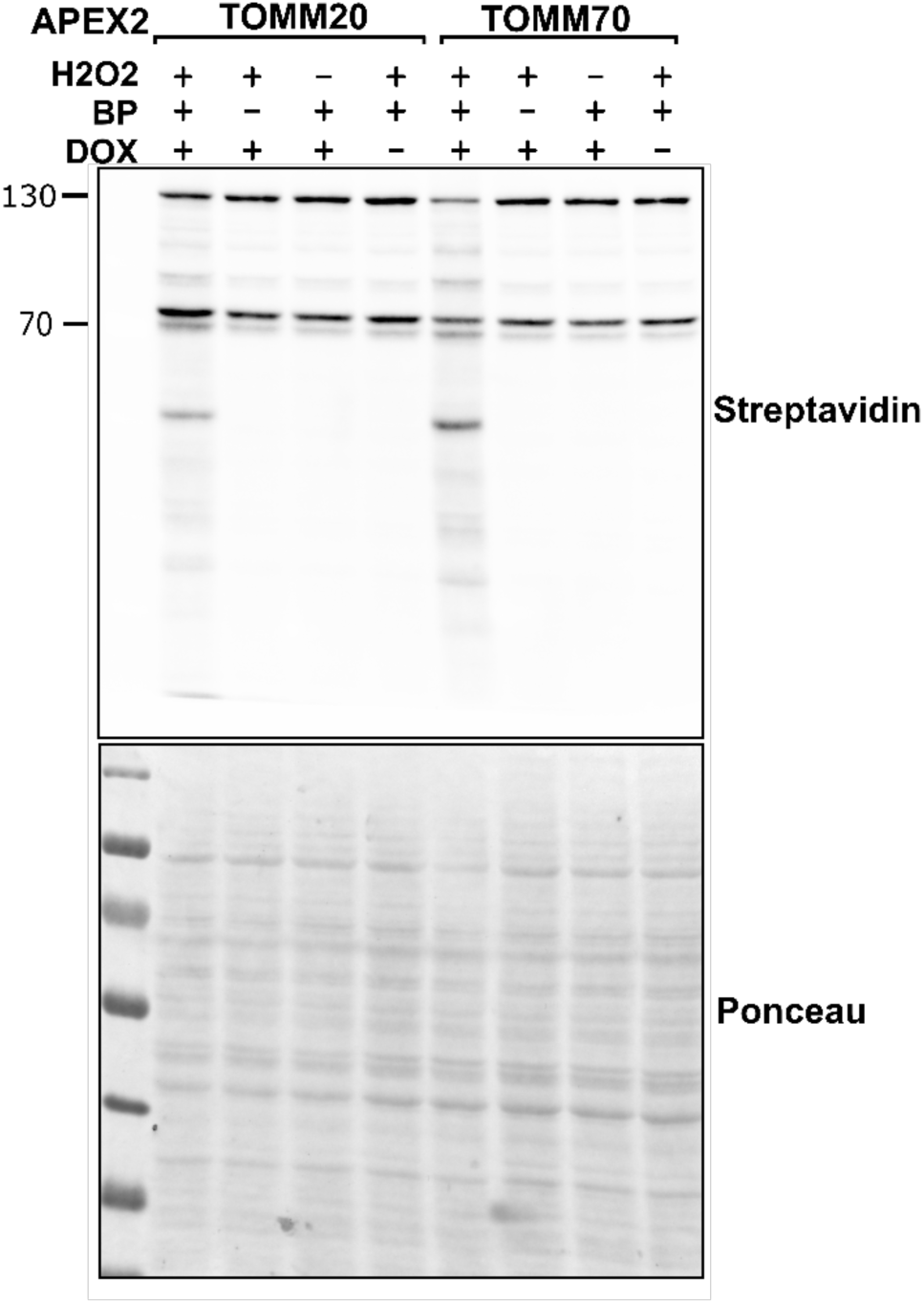
Target protein biotinylation is mediated by TOMM20-or TOMM70-APEX2 fusion proteins. Western blot analysis of cell lysates to confirm the APEX2-mediated biotinylation in Hela cells stably expressing the indicated APEX2 fusion proteins. The blot is stained either with Streptavidin-HRP conjugate to detect biotinylated proteins (upper part) or with Ponceau (lower part). Biotinylation depends on the presence of both biotin phenol (BP) and H_2_O_2_. Omission of either substate or lack of APEX2 fusion protein expression only results in detection of endogenous biotin-containing proteins at ∼75 and ∼150 kDa.

**Figure 3.**
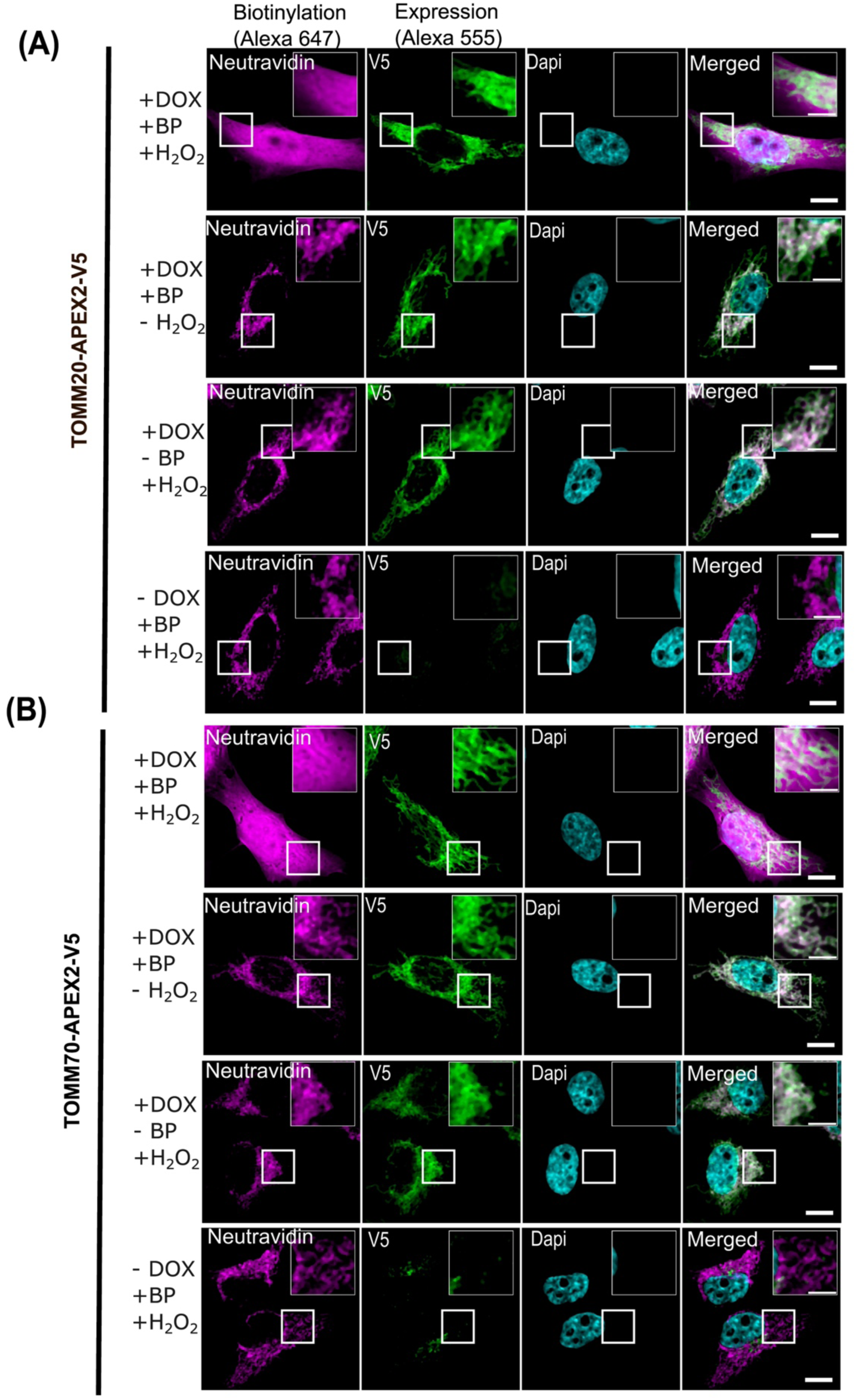
Local biotinylation at mitochondria by TOMM20– and TOMM70-APEX2. Confocal fluorescence imaging of APEX2-mediated biotinylation in Hela cells stably expressing the indicated APEX2 fusion proteins. (**A**) TOMM20-APEX2-V5 (**B**) TOMM70-APEX2-V5. After 24 h of DOX induction, cells are exposed to live-cell biotinylation with Biotin phenol (BP) and H₂O₂ for one minute and subsequently fixed. Cells are immunolabeled with ‘V5’ to check the expression of indicated APEX2 fusion proteins (green) and ‘Neutravidin’ to stain the biotinylated species (Magenta). When either H₂O₂ or BP is omitted, or when the cells are not induced, the endogenous mitochondrial biotinylated proteins become apparent. Nuclei are stained with DAPI (cyan). Incents are the magnified portion of the cell showing on the uppermost right of each panel (Scale bar: 10 μm; incent: 5 μm).

### LFQ-based quantitative proteomics approach to study TOMM20– and TOMM70 interactome

Since the TOMM20– and TOMM70-APEX2 fusion proteins are expressed at endogenous levels and correctly associated with endogenous TOM complex proteins, we next aimed to identify their corresponding interactomes. To control for specific interactions, we transfected and expressed two additional APEX2 fusion proteins as controls into HeLa-EM2-11ht (**Suppl. Figure 1A**) ^29,33^. Mito-APEX2 targets the APEX2 enzyme to the mitochondrial matrix. A second control protein, NES-APEX2 targets APEX2 to the cytosol due to a nuclear export signal (NES)^29^. Like the TOMM20– and TOMM70-APEX2 fusions, the two constructs are expressed by addition of doxycycline(‘DOX’) (**Suppl. Figure 1B**). As described before^29,33^, Mito-APEX2 is targeted to the mitochondria (**Suppl. Figure 1C**) as shown by co-localization with the endogenous marker protein TOMM22. The diffuse intracellular staining pattern and nuclear exclusion of NES-APEX2 demonstrates its cytosolic location (**Suppl. Fig. 1C**). Both control APEX2 proteins are functional and increase detectable biotinylation only in the presence of BP and H₂O₂ (**Suppl. Figure 2A**). In case of Mito-APEX2, neutravidin-Alexa647 staining of biotinylated proteins *in-situ* (**Suppl. Figure 2B**) shows a very similar distribution to that of the enzyme itself, most likely due to the confinement of the activated phenoxyl-biotin radicals in the mitochondrial matrix. As expected, the distribution of proteins biotinylated by NES-APEX2 is more diffuse and mimics the distribution of cytoplasmic proteins.

Six independent replicates for TOMM20-APEX2, and three for TOMM70-APEX2, NES-APEX2, and Mito-APEX2 were analyzed by bottom-up proteomics. In addition, we included three replicates of control experiments performed with cells not expressing the corresponding TOM complex resident bait proteins (‘-DOX’). Biotinylation with BP and H₂O₂, quenching, lysis and capturing was essentially done according to previously published protocol^34^ (see Methods). The captured proteins were further measured by liquid chromatography-tandem mass spectrometry (LC-MS) after on-bead tryptic digestion. Downstream data processing was performed as label free quantification (LFQ), and imputation of missing values was applied during subsequent data analysis to increase the number of identifications. With this approach we identified in total 2,177 protein groups of which 499 are annotated for mitochondrial localization by MitoCarta3.0^35^. Except for –DOX controls, 1,300 to 1,700 protein groups were identified with the highest number of mitochondrial localized proteins for Mito-APEX2 (**Suppl. Figure 3A**). Overall, we observed an excellent correlation between the replicates (**Suppl. Figure 3B**). We next evaluated the sub-organelle distribution of proteins identified in TOMM20– and TOMM70-APEX2 interactomes (**Suppl. Figure 3C**). Around 1700 proteins were identified in TOMM20-APEX2 and TOMM70-APEX2 indicating a very similar number of biotinylated proteins. Besides the expected mitochondrial proteins, we also surprisingly observed enrichment of nuclear proteins, ribosomal, and cytosolic proteins in the interactomes of both bait proteins. While the appearance of nuclear proteins is unclear but has already been observed in the interactome of a TOMM20-TurboID^36^ protein fusion, cytoplasmic proteins in TOMM20– and TOMM70-APEX2 interactomes might be due to the positioning of the APEX2. Since the APEX2 parts of the two fusion proteins are facing towards the cytosol, we expected them to not only biotinylate MOM proteins but also, due to the release and diffusion of the phenoxyl-biotin radicals, proteins in the surrounding cytoplasm^32^.

To validate that the TOMM20– and TOMM70-APEX2 bait constructs allow us to study their respective interactomes, we compared the biotinylated proteomes generated by these two baits with those of cells not induced for expression of the corresponding APEX2 constructs (**Figure 4 A and B**; **Suppl. table 1 and 2)**. As expected, much fewer proteins were identified in the uninduced controls, making imputation of especially low abundant proteins for these samples mandatory (**Suppl. Figure 3 D-G**). Among the identified candidates, we further distinguished significantly abundant ones (p value ≤ 0.05) and stringently-significantly abundant ones (p ≤ 0.01). The identification of several other subunits of the TOM complex (e.g. TOMM40) or other MOM proteins (21 in case of TOMM20-APEX2, seven in case of TOMM70-APEX2) among the significantly and stringent significantly high abundant proteins demonstrates the effectiveness of our approach.

**Figure 4.**
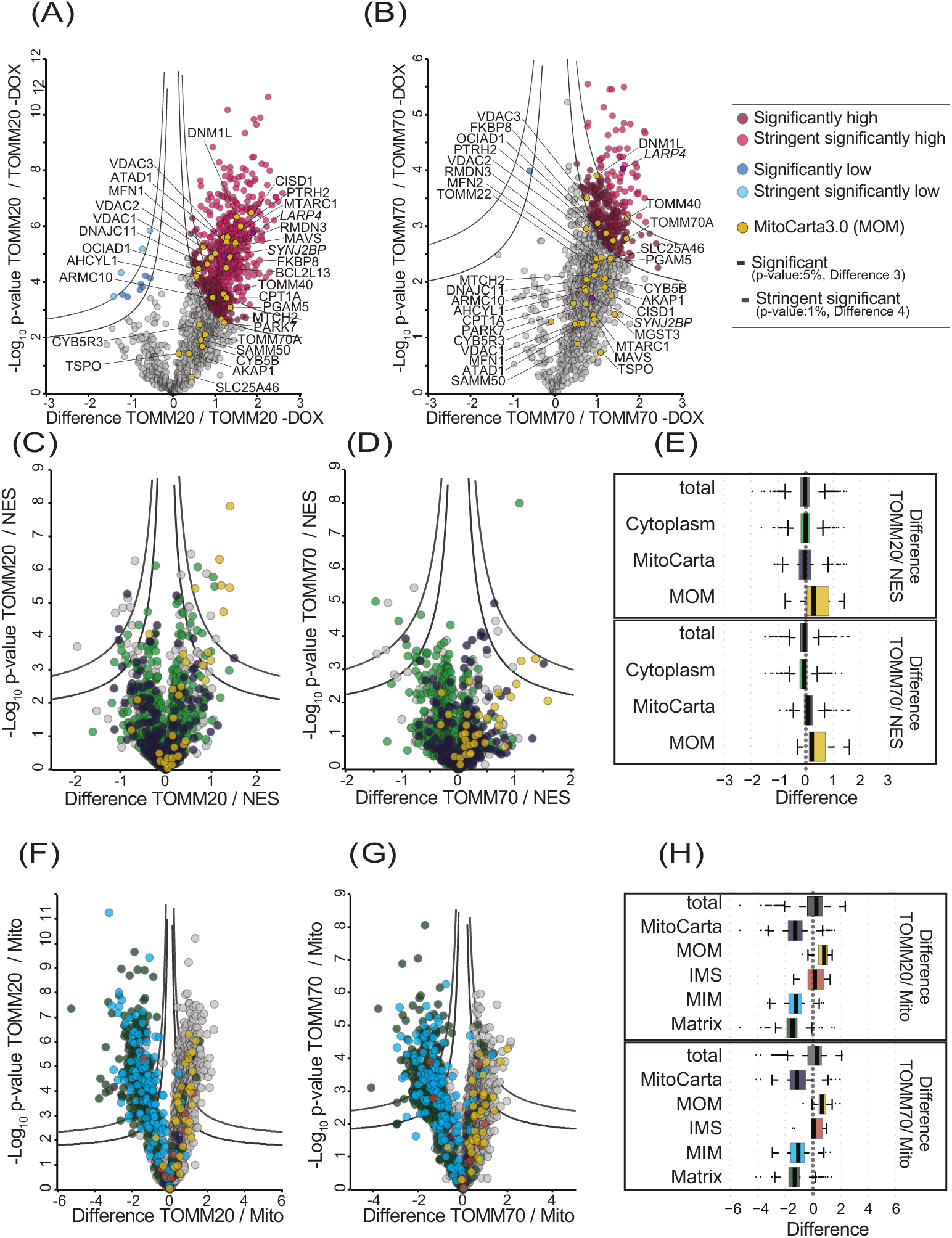
Interactomes of TOMM20-APEX2 and TOMM20-APEX2 at the mitochondrial outer membrane. Interactome of TOMM20-APEX2 **(A)** and TOMM20-APEX2 **(B)** including many mitochondrial proteins compared to their respective uninduced (-DOX) controls. Volcano plots for TOMM20-APEX2 **(C)** and TOMM70-APEX2 **(D)** against NES-APEX2 interactome. Highlighted are proteins annotated for cytoplasmic localization (based on GOCC) and mitochondrial localization (MitoCarta3.0). The color code used to annotate the proteins in C and D is identical to E (**E)** Boxplot with distribution of total proteins compared to proteins annotated for mitochondrial, and cytosolic localization for TOMM20-(above) and TOMM70-APEX2 (below) against NES-APEX2. Volcano plots for TOMM20-APEX2 **(F)** and TOMM70-APEX2 **(G)** against Mito-APEX2 interactome. Highlighted are proteins annotated for their respective submitochondrial localization based on MitoCarta3.0. The color code used to annotate the proteins in F and G is identical to H. **(H)** Boxplot with distribution of total proteins compared to proteins annotated for mitochondrial, and mitochondrial sub localization for TOMM20-(above) and TOMM70-APEX2 (below) against Mito-APEX2. Indicated are thresholds for stringent significantly higher or lower abundant proteins (p-value 1%, Difference 4) and significantly abundant proteins (p-value 5%, Difference 3).

Surprisingly, whereas TOMM70 was present in the dataset of potential interactors of TOMM20-APEX2 and identified as a biotinylated protein in TOMM70-APEX2 expressing cells, TOMM20 was not. A closer look at the raw data revealed that it had been identified only by one peptide, and due to the raw data processing settings, it was not considered for further analysis. Possible reasons for the low number of TOMM20 peptides are the low number of the peptides that are generated from the rather small 20 kDa protein itself and the limited accessibility of the rather small cytosol-facing domain of TOMM20^37^ to activated biotin. In addition, the cytoplasmic domain contains few aromatic amino acids available for reaction with the biotin-phenoxyl radical generated by APEX2^34,37^.

### TOMM20-/ TOMM70-APEX2 interactome at the MOM-Cytoplasm interface

Next, we validated that the interactome of TOMM20– and TOMM70-APEX2 indeed reflects processes occurring at the MOM-Cytoplasm interface. We compared the pattern of biotinylated proteins with that of a cytosolically targeted APEX2 (NES-APEX2). Importantly, a first observation made when comparing the interactome of TOMM20– and TOMM70-APEX2 with that of NES-APEX2 was that MOM proteins were more abundant among the TOMM20– and TOMM70-APEX2 interactomes (**Figure 4 C-E**; **Suppl. Table 3 and 4).** However, cytoplasmic proteins were also abundant in both differential interactomes (TOMM20-APEX vs. NEX-APEX and TOMM70-APEX2 vs NES-APEX2; see also Supplemental Figure 3C). As one of our aims was to elucidate the extent of RBPs and factors involved in (local) translation that are associated with TOMM20 or TOMM70, we next analyzed our dataset for the enrichment of proteins related to RNA function. While no RBPs were identified as enriched in the TOMM70-APEX2 interactome when compared to that of a cytoplasmic APEX2, seven RBPs (i.e. 23% of all enriched proteins compared the NES-APEX2) were found in the TOMM20-APEX2 interactome, including SYNJ2BP, PAIP1, MAVS, and PABPC4L (**Suppl. table 3**). To verify that these proteins can indeed be found in the proximity of the TOM complex, we performed super-resolution microscopy, using antibodies against TOMM20 and SYNJBP2 (**Suppl. Figure 4**). Quantification of the signals revealed a high positional overlap of both proteins, supporting the proximity labeling data.

Next, we compared the interactome of TOMM20– and TOMM70-APEX2 against that of a matrix-targeted APEX2 (Mito-APEX2; **Figure 4 F-H**; **Suppl. table 5 and 6**). However, in comparison to Mito-APEX2 both MTS receptor APEX2 baits showed higher abundance of annotated MOM proteins and low abundance of matrix or MIM proteins. This validates that our APEX2-based approach mainly targets proteins localized to the MOM or to the MOM-Cytoplasm interface in close proximity to the TOM complex (**Figure 4 F-H**). Interestingly, in addition to several MOM proteins, we identified additional mitochondrial proteins of the matrix, MIM or IMS. Compared to TOMM70-APEX2, for TOMM20-APEX2 we identified more MOM annotated proteins (14 as compared to six). Besides the MOM proteins, additional interactors of TOMM70 belong to IMS, MIM, and matrix protein groups. Overall, a comparison of significantly enriched candidates in TOMM20 or TOMM70 vs Mito-APEX2 revealed 17 proteins enriched only in TOMM20 vs Mito-APEX2, 12 proteins in TOMM70 vs Mito-APEX2, and five overlapping proteins in both interactomes (**Suppl. Figure 5A)**. Additionally, TOMM20 appears to interact with more MOM proteins compared to TOMM70, probably due to a more stable association with the MOM or the TOM complex^7–9^.

After we validated the suitability of our experimental design to study the interactome of TOMM20– and TOMM70-APEX2 at high resolution at the interface of MOM and cytoplasm, we next compared the differential interactomes of TOMM20– and TOMM70-APEX2 (**Figure 5A**). While most mitochondrial proteins are similarly abundant between the two baits, 20 proteins were identified to be significantly overrepresented in the TOMM70-APEX2 interactome including three proteins annotated as exclusively mitochondrial; NDUFA4(MIM), PPA2 (matrix), and seven proteins annotated as dual or multiple localized including pyruvate carboxylase (matrix, cytosol), MCCC1 (matrix, cytosol), UQCRC2 (MIM, nucleoplasm), MFN2 (MOM, ER, cytosol), GPX4 (MIM, extracellular space, nuclear envelope, cytosol), MACROD1 (matrix, cytoplasm, nucleus), and SRRYD4 (cytosol, matrix) (**Suppl. table 7**). A total of 35 proteins were found to be significantly overrepresented in the TOMM20-APEX2 interactome, among them four proteins annotated as mitochondrial including DLAT (Matrix), TST (Matrix, extracellular matrix), TXNRD2 (Matrix, cytosol), and DNAJA1 (MOM, ER, cytosol, nucleus, extracellular matrix) (**Figure 5A**; **Suppl. table 8**). We cannot prove without doubt if these proteins are mature mitochondrial proteins or precursors that were caught while interacting with the TOMM proteins. Likely due to the overall low coverage of peptides in the MTS regions, hardly any MTS containing peptides expected to be part of the precursors were identified. Thus, distinguishing between precursors in transit and mature proteins already at their destination site was not possible. However, previous studies showed that biotin-phenoxyl radicals generated by APEX2 at the cytoplasmic face of mitochondria may occasionally traverse the MOM through porins and can biotinylate MIM and IMS proteins. However, these radicals cannot penetrate the matrix due to the selective permeability of MIM ^34,38^. This suggests that the biotinylated matrix proteins in the TOMM20-APEX2 interactome are captured rather during transit than after processing in the matrix. Our results thus support the idea that mammalian TOMM20 and TOMM70 receptors, like their yeast counterparts^2,3^ might interact with distinct but overlapping sets of precursor proteins.

**Figure 5.**
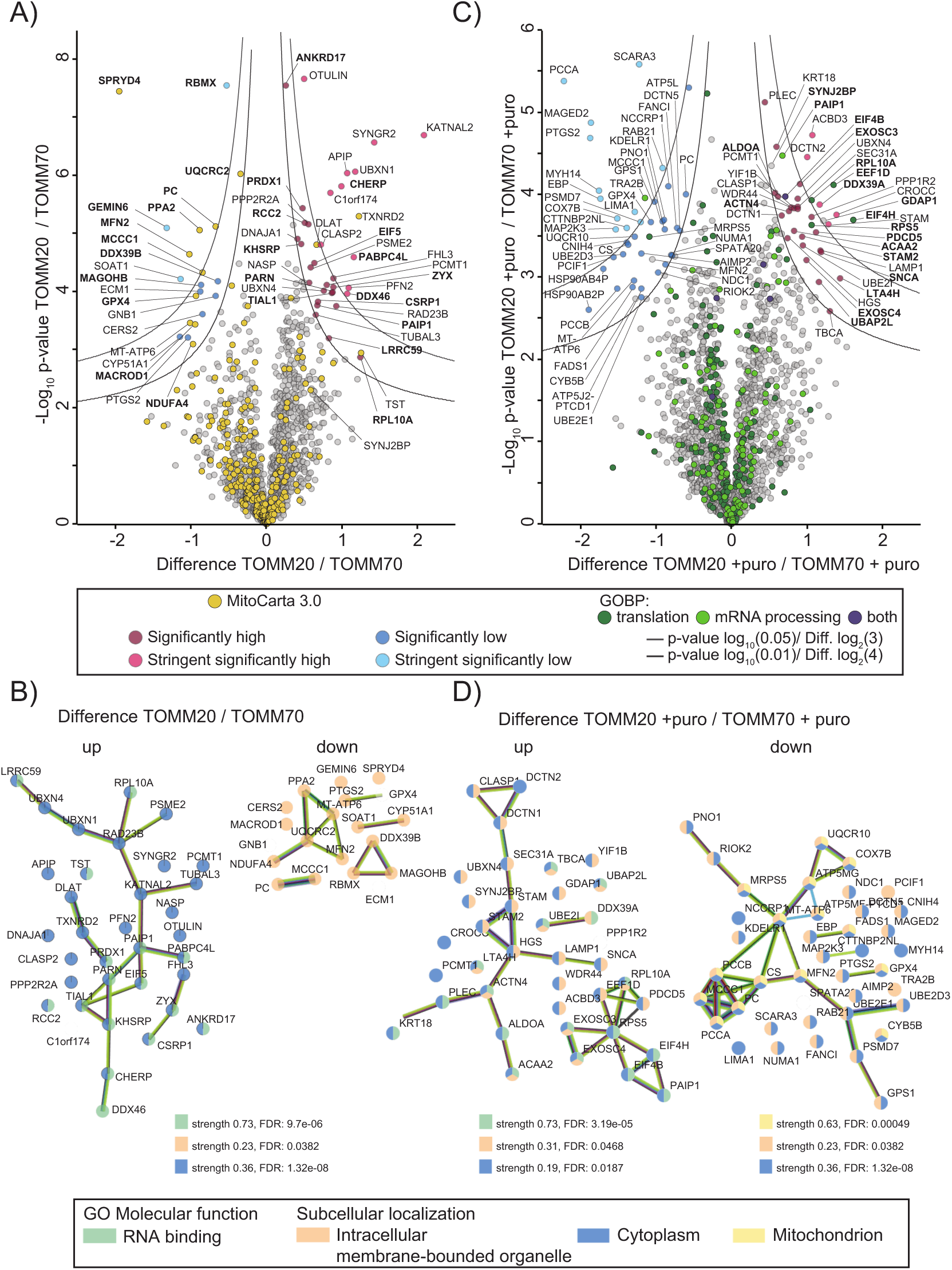
Differential interactome of TOMM20-APEX2 versus TOMM70-APEX2. (**A**) Volcano plot for TOMM20-APEX2 interactome against TOMM70-APEX2 interactome. Yellow highlighted proteins are annotated for mitochondrial localization (MitoCarta3.0). Indicated are thresholds for stringent significantly higher or lower abundant proteins (p-value 1%, Difference 4) and significantly abundant proteins (p-value 5%, Difference 3). **(B)** STRING analysis of proteins abundant in differential interactome of TOMM20-APEX2 vs TOMM70-APEX2 from volcano plot A, showing enriched candidates of TOMM20 (up), and enriched candidates of TOMM70 (down). Proteins are annotated according to GO molecular function and subcellular localization. **(C)** Volcano plots for TOMM20-APEX2 interactome under puromycin treatment (+puro) against TOMM70-APEX2 interactome (+puro). Highlighted are the proteins annotated (based on GOBP) for translation and mRNA processing **(D).** STRING analysis of proteins abundant in the differential interactome of TOMM20-APEX2 (+ puro) vs TOMM70-APEX2 (+puro) from volcano plot C showing enriched candidates of TOMM20 (up), and enriched candidates of TOMM70 (down). Proteins are highlighted according to GO molecular function and subcellular localization. Strength and False discovery rate (FDR) of each pathway are indicated.

### RNA-binding proteins and translation factors are part of the TOMM20-APEX2 interactome

A surprisingly large number of cytoplasmic RBPs including CHERP, PAIP1, KHSRP/FUBP2, PABPC4L, TIAL1, PARN, DDX46, ANKRD17, LRRC59, and components of the translational machinery like RPL10A and EIF5 were identified as significantly enriched in TOMM20-APEX2 compared to TOMM70-APEX2. To investigate this further, we subjected the significantly enriched proteins in the differential TOMM20 vs TOMM70 interactome to a STRING-based network analysis^39^.The majority (43%) of these proteins in the TOMM20-APEX2 interactome are overrepresented for cytoplasmic localization with multiple of them annotated as RBP (**Fig 5 B; Up**). STRING analysis further reveals that most of the processes the enriched RBPs are involved in are interconnected, supporting our hypothesis that TOMM20-APEX2 specifically biotinylates a distinct and functionally related group of RBPs and translation factors that are in its close proximity, rather than doing so randomly. We further checked if these proteins were also abundant in our other comparisons of interactomes (TOMM20-APEX2 vs –DOX, TOMM20-APEX2 vs NES-APEX2, and TOMM20-APEX2 vs Mito-APEX2; **Suppl. Table 9**), which revealed that many RBPs and translation factors including PAIP1, PABPCL, and EIF5 were enriched in the TOMM20 interactome versus these other ones. In contrast, the STRING analysis of the TOMM70-APEX2 revealed that most of the proteins that are more enriched in its interactome versus that of TOMM20-APEX2 are linked to membrane bound organelles, including one cluster consisting of mitochondrial proteins (PPA2, UQCRC2, MFN2, and NDUFA4; **Fig 5B; down**). We analyzed the interaction profile of the TOMM20 interactome derived from the comparison of TOMM20 vs. TOMM70 (35 proteins, see figure 5A) with the enriched proteins in TOMM20 vs NES-APEX2 interactome (31 proteins: see figure 4C). Most of the higher abundant proteins (36%) shared between these interactomes are annotated as either RNA binding or translation (**Suppl. Figure 5B**).

The identification of several RBPs or proteins involved in translation in our enriched proteomes tempted us to investigate how the interactomes of TOMM20– and TOMM70-APEX2 changes in the presence of the translation inhibitor puromycin. For that, we included three replicates of puromycin treated (+puro) samples for TOMM20– and TOMM70-APEX2 expressing cells. For treatment, we chose a 30 min window and 200 μM concentration of puromycin since a similar treatment had revealed only little change in the overall proteome in HEK293 cells^16,27^. Puromycin treatment (+puro) of TOMM20-APEX2 expressing cells resulted in surprisingly little change in the associated proteome with only one protein significantly higher abundant (PRTEDC1) or lower abundant (FANC1, CHERP, and FADS1) after treatment **(Suppl. Figure 5C; Suppl. Table 10)**. However, none of these proteins have an obvious connection to each other, to RNA, or to mitochondrial function. A similar observation was made upon treatment of TOMM70-APEX2 cells with puromycin although the number of lower or higher abundant proteins was slightly larger (**Suppl. Figure 5D; Suppl. Table 11**). However, when we compared the differential interactome of TOMM20 vs TOMM70 (both after treatment with puromycin) and focused especially on proteins annotated by GOBP as being involved in translation and mRNA processing, we found in total 258 proteins, of which eleven were significantly more abundant in the TOMM20-APEX2 after puromycin treatment vs TOMM70-APEX after exposure to puromycin (**Figure 5C; Suppl. Table 12**). These include the translation initiation and elongation factors (eIF4B, eIF4H, eEF1D), ribosomal proteins (RPS5, RPL10A), and RBPs that are primarily involved in controlling mRNA stability. Among the latter proteins are PAIP1, a translational co-activator interacting with polyA-binding protein PABP; SYNJ2BP, a MOM-localized RBP that binds to mRNAs encoding mitochondrial proteins, mitigates the effects of translation stress, and has been linked to the local translation at the mitochondrial surface^27^; and finally SNCA, an RBP that forms amphipathic helix, similar to the MTS motifs^40^, and an known interactor of TOMM20^41,42^.The increased abundance of RBPs that are involved in the mRNA stability in the TOMM20-APEX2 dataset after translation inhibition suggests that TOMM20, via interacting with these proteins has a more mRNA-protective role than TOMM70. Though depletion of these candidates from TOMM70 differential interactome after puromycin treatment suggests that TOMM70 is also dynamically involved in their translation-dependent mitochondrial targeting.

In a similar analysis as before (Figure 5B), the significantly enriched proteins in the TOMM20-APEX2 (+puro) (**Figure 5D; up**) and TOMM70-APEX2 (+puro) (**Figure 5D; down**) datasets were subjected for STRING-based network analysis. For the TOMM20-APEX2 interactome, we identified a cluster of proteins annotated as translation regulators. This group of proteins includes a RBP (PAIP1), translation factors (eIF4B, eIF4H, EEF1D), and ribosomal protein (RPS4), suggesting that components of the translation machinery are still proximal to TOMM20 upon puromycin inhibition.

Interestingly multiple proteins were enriched for their mitochondrial inner membrane and matrix localization among the significantly high abundant interactors of TOMM70-APEX2 after puromycin treatment (**Figure 5C; Suppl. Table 13)**. Specifically mitochondrial matrix proteins such as MCCC1 (encoding a subunit of the 3-methylcrotonyl-CoA carboxylase) and PC (pyruvate carboxylase) are similarly overrepresented even under non-translation inhibitory conditions, suggesting a translation-independent import of these proteins.

In summary, our quantitative proteomic approach using differential interactomes supports the hypothesis that the TOMM20 and TOMM70 receptors interact with unique sets of proteins. More importantly, our analysis provides first evidence that both receptors remodel their proteome differentially in response to the translation stress.

## Discussion

Proximity labeling approaches have been extensively used in the past to investigate the mitochondrial proximal proteome and transcriptome^16,27,34,43^. In most of these studies, the proximity ligase was targeted to the mitochondrial outer membrane by fusing it to a targeting peptide from the MAVS protein^27,34^. Although this approach allows MOM targeting of APEX2 or TurboID enzymes, it does not reveal if the identified associated proteins are in a functional relation to the TOM complex that controls mitochondrial protein import. To address this question more directly, we used proximity labeling based on APEX2 and quantitative mass spectrometry to investigate the presence of mRNA-interacting proteins in the proximity of the TOM complex at the MOM and compared the interactomes of mammalian TOMM20 and TOMM70 in HeLa cells.

We validated our approach by demonstrating an enrichment of MOM proteins in proximity to the TOM complex, especially when comparing TOMM-APEX2 interactomes with that of cytoplasmic or mitochondrial matrix localized APEX2. Particularly in case of TOMM20-APEX2, we identified, besides expected MOM components like MTARC1 (a MOM-localized oxidoreductase), OCIAD1 (OCIA domain-containing protein 1), CISD1 (a redox active mitochondrial protein with an iron–sulfur domain), RMDN3 (a regulator of microtubule dynamics), BCL2L13 (a BCL2-like protein), and CYB5R3 (NADH-cytochrome b5 reductase 3), RNA-binding proteins. These include SYNJ2BP and MAVS that had already previously been linked to the MOM^27,44^.

In a related study, the interactome of human TOMM20 was recently captured in HCT116 cells by tagging it with the miniTurbo biotin ligase^36^. Around 22% proteins significantly abundant in our differential dataset of our TOMM20-vs NES-APEX2 dataset were also enriched in their interactome. These proteins include MOM proteins like CISD1, RMDN3, MTARC1, OCID1, and MAVS but only eIF5 as translation-related protein. The low overlap of enriched proteins between the studies might be due to the different expression setups, proximity labeling tags and labeling conditions as it has been observed e.g. for the TDP-43 protein^45^. Since transient interactions at MOM are crucial to modulate mitochondrial quality control and function, our approach is beneficial in capturing the one-minute snapshot of such transient and spatial interactions of both TOM receptors.

The direct comparison of TOMM20 vs TOMM70 confirmed that both receptors, though belonging to the same TOM complex, interact with different substrates of mitochondrial preproteins destined for various submitochondrial compartments as seen for their yeast homologs^46^. When focusing on the mitochondrial proteins associated with TOMM20-APEX2, we also saw that these mainly belong to matrix proteins while those enriched in the TOMM70-APEX2 interactome represent matrix and mitochondrial inner membrane (MIM) proteins. This supports previous studies showing that MTS-containing matrix precursor proteins are mainly imported by TOMM20^47^ or that the loss of Tom70/71 in yeast also in particular affects import of matrix and MIM proteins^46,48^. Furthermore, our data also indicates that the receptors are in proximity to a unique set of proteins at the MOM, as shown by a significant difference among their MOM-specific interactomes.

TOMM20-in contrast to TOMM70-APEX2 appears to associate with cytoplasmic RBPs like PAIP1, KSHRP, and PABPC4L, and components of translation machinery players like ribosomal proteins or the translation initiation factor EIF5. This enrichment of cytoplasmic RBPs and components of the translation machinery not only corroborates the hypothesis of localized translation at the MOM^18,22,23^ but also suggests that TOMM20 rather than TOMM70 might play a role in localized translation of nuclear encoded mitochondrial mRNAs or co-translational import of their encoded proteins.

The remodeling of its interactome during inhibition of translation suggests that TOMM20 might play a role in preserving cellular hemostasis during translation stress by retaining translation-related proteins at the MOM. It specifically interacts with the RPBs involved in mRNA stability like SYNJ2BP, SNCA, or PAIP1. SYNJ2BP has previously been identified as a key regulator to safeguard specific nuclear-encoded mitochondrial transcripts during translation stress^27^. It anchors these transcripts under stress recovery which might facilitate their local translation and import, thereby maintaining OXPHOS activity and mitochondrial function^27^. We captured not only this RBP in the TOMM20 interactome under translation arrest but also other components of the translation machinery like eiF4B, eiF4H, eEF1D, or ribosomal proteins, providing more evidence for a SYNJ2BP mediated localized translation at the MOM in proximity of the TOMM20 receptor. PAIP1 interacts with key translation factors and modulates the stability and translation of mRNAs^49–51^. SNCA is a known interactor of TOMM20^42^, and TOMM20 overexpression reportedly rescued α-Synuclein induced dopaminergic neurodegeneration in Parkinson’s disease patients^41^. The presence of these RBPs in TOMM20 interactome under translation inhibition conditions could reflect their direct role as safe guarders against translation stress, which still needs to be investigated. In the future, the integration of similar APEX2-based proximity labeling approaches of other TOM complex subunits or TOM-associated proteins like the SAM complex^52^ with the presented data could reveal a more detailed view of the complex interactions at the MOM and provide further insights into the processes of localized translation at the MOM and its contribution to mitochondrial import.

## Material and Methods

### Cell culture

Each APEX2 fusion protein expressing cell line was generated by using a low passage HeLa 11ht parental cell line obtained from Dr. Kai Schönig (Zentralinstitut für Seelische Gesundheit Mannheim, Germany)^53^. Cells were always cultured in DMEM (Sigma) mixed with 10% fetal bovine serum (FBS), 110 mg ml^−1^ Sodium Pyruvate, and 1x Penicillin/Streptomycin at 37°C with 5% CO2, and maintained by 200 μg ml^−1^ Hygromycin B (Sigma) and 200 μg ml^−1^ G418 (Sigma) as described before ^54^.

### Plasmid construction

All plasmids were generated via Gibson assembly. As a backbone for all constructs, plasmid RJP2501 (a pSF3 backbone vector containing GFP) was digested with restriction endonucleases BglII and PacI to excise the GFP gene. This resulting linearized vector was used as a template to generate all plasmids via Gibson assembly (Supplementary table 14). Each corresponding fragment of the insert contains overlapping ends by the respective primers to facilitate the seamless joining of adjacent fragments via Gibson assembly reaction.

### Generation of stable cell lines

To enable tunable doxycycline (DOX) induced expression and preserve isogenicity, the fusion protein expression cassette in the generated pSF3 plasmid for each cell line was stably integrated in the genome of the Hela 11ht cells via recombinase-mediated cassette exchange (RMCE) as described before^54^. Each doxycycline inducible cell line expressing APEX2 containing fusion protein was obtained by selection with 50 μM ganciclovir (Sigma).

### Immunofluorescence microscopy

Around 25,000 cells were seeded on the coverslips coated with 0.2% Gelatin placed in 12-well plates. Cells were incubated either with or without 500 ng ml^−1^ doxycyclin for 24 hours. The samples were then thoroughly washed three times with PBS to remove media, fixed and permeabilized with ice-cold methanol for 10 minutes at –20°C. The samples were washed two times with PBS containing 5 mM MgCl_2_ (PBSM) and once with 50 mM Glycine in PBSM. Cells were further incubated in 3% bovine serum albumin (BSA) diluted in PBSM (blocking solution) for 1 h. This was followed by incubation with the corresponding primary antibodies (mouse anti-V5 antibody,1:1000 dilution; rabbit anti-TOM22 antibody,1:200 dilution; mouse anti-TOM20 antibody,1:200 dilution; rat anti-flag antibody,1:500 dilution; rabbit anti-TOM20 antibody, 1:200 dilution; further information on antibodies can be found in supplementary table 15) in the blocking solution for 1 h at room temperature or overnight at 4 °C. After washing three times with PBS, cells were incubated with Alexa Fluor coupled secondary antibodies (Invitrogen) at 1:1000 dilution in the same blocking buffer for 1 h. The cells were washed two times with PBS first and then incubated for 5 minutes with DAPI (0.1 μg ml^−1^) diluted in PBS. Cells were thoroughly washed again with PBS and mounted with ProLong™ Glass Antifade Mountant (Thermo). After overnight incubation at room temperature, the slides were stored at 4 °C until imaging. To analyze biotinylated proteins, biotinylation was performed directly on the cells seeded on the cover slips as explained below before treatment with the corresponding antibodies and mounted as explained before. All images were captured using a Zeiss LSM980 laser scanning confocal microscope (Carl Zeiss, Germany) equipped with an Airyscan 2 detector with a 60x oil objective and further processed with ImageJ/Fiji.

### Cell fractionation

For each cell fractionation experiment, cells were induced with 500 ng ml^−1^ doxycycline for 24 hours or mock treated. Following induction, media was discarded from the around 85% confluent cell culture dishes, and cells were washed with PBS twice and then collected in 15 ml centrifuge tubes. Cells were pelleted at 500 × *g*, 3 min at 4°C, the supernatant discarded, and the cell pellet resuspended in HMS-A buffer^55^ (0.22 M mannitol, 0.02 M HEPES-KOH, pH 7.6, 1 mM EDTA, 0.075 M sucrose, 0.1% BSA, 1X cOmplete™protease inhibitor cocktail (Roche)). The suspension was incubated for 10 minutes on ice, transferred to a homogenizer (Sigma) and lysed with 30 cycles on ice. The lysates were centrifuged at 900 × *g*, 5 min at 4°C. The supernatant was collected and centrifuged again for 9000 × *g*, 15 min at 4 °C, resulting in a crude mitochondrial fraction in the pellet and a cytosolic fraction in the supernatant. The mitochondrial pellet was further washed with HMS-B buffer (HMS-A buffer without BSA) and mitochondria pelleted again at 10,000 × g, 10 min, 4 °C. To solubilize mitochondria, the pellet was resuspended again in HMS-B buffer with digitonin (4:1 (w/w) ratio of detergent to the protein) and incubated for 30 minutes at 4°C. Non-solubilized material was removed via centrifugation (13,000 × *g*, 10 min, 4 °C). The solubilized mitochondrial fraction was collected and denatured for Western blot. Around 100 µg solubilized mitochondrial fraction was separated either on a 12% SDS-PAGE gel (for TOMM20-APEX2 detection) or a 10% SDS-PAGE gel (for TOMM70-APEX2 detection). For expression validation in Figure 1F, blots were incubated with primary antibodies 1 hour at room temperature or overnight at 4°C. Primary antibodies include mouse anti-TOM20 (1: 250 dilution), rabbit anti-TOM40 (1: 500 dilution); mouse anti-TOM70 (1:500 dilution), and mouse anti-GAPDH (1:1000 dilution).

### Co-immunoprecipitation

Mitochondria were isolated and solubilized as described above. Around 250 µg of solubilized mitochondria were used for each co-purification and incubated with 10 µg of anti-TOM22 (Abcam) or 20 µg of anti-TOM40 (Proteintech) antibodies for 2 h at 4°C. Normal rabbit IgG was used as an isotype control. After incubation, prewashed protein A agarose beads were added and incubated for another hour at room temperature. Beads were washed three times with HMS-B buffer and proteins eluted with 2 × Laemmli buffer at 95°C for 10 min. For V5 co-immunoprecipitation, V5 beads (ChromoTek) were pre-washed with HMS-B buffer three times. Solubilized mitochondria were crosslinked first with 1mM DSP for 1 h at 4°C followed by quenching using 1 M Tris-HCl, pH 7.5. The crosslinked solubilized mitochondria were purified through a filter column and incubated with 50 µl of V5 beads in HMS-B for 1h at room temperature. Beads were washed three times with HMS-B buffer, eluted in 2X Laemmli buffer with 0.1 M DTT at 95°C for 10 min, and further analyzed by Western blot. After boiling the beads in the Laemmli buffer, the eluate was collected, separated on a 12% SDS-PAGE gel before detection of proteins by Western blotting. The primary antibodies used were rabbit anti-APEX2 (1:3000 dilution), rabbit anti-TOM22 (1:500 dilution), and rabbit anti-tomm40 (1:500 dilution), and mouse anti-V5 (1:1000 dilution).

### Proximity labeling

For APEX2 labeling, cells were induced with doxycycline (DOX) or mock treated as described above. After 24 h, cells at around 85% confluency were incubated with fresh DMEM containing 0.5 mM biotin-phenol (BP; Iris Biotech GMBH) for 30 min at 37°C. Biotinylation was initiated by adding H_2_O_2_ for one minute at room temperature to the final concentration of 1 mM under constant agitation for each cell line^34^. Control samples omitted either BP or H_2_O_2_ or uninduced cells were included for each APEX2-fusion protein expressing cell lines. Labeling reactions were quenched by removing media and immediate washing of cells with an equal volume of quenching DPBS solution that was prepared freshly after mixing DPBS with 10 mM sodium azide, 10 mM sodium ascorbate, and 5mM Trolox. Cells were further washed two times with quenching DBPS solution, and once with PBS before trypsinization, and pelleting at 500 × *g*. The pellet was washed with PBS and immediately lysed in RIPA buffer (50 mM Tris, pH 7.5, 150 mM NaCl, 0.1% SDS, 1% Triton X-100, 0.5% sodium deoxycholate, and 1x cOmplete™protease inhibitor cocktail) supplemented with 10 mM sodium azide, 10 mM sodium ascorbate, and 5 mM Trolox. The lysates were incubated on ice for 15 minutes, sonicated briefly, then centrifuged at 15,000 × *g* for 10 min at 4°C. Protein concentration of the clarified lysates was determined with a Pierce^TM^ 660-nm protein assay (Thermo, catalog no. 22660). 20 µg of whole cell lysate was combined with Laemmli buffer and boiled for 10 min. Lysates were separated on a 9% SDS-PAGE gel and transferred to nitrocellulose membrane. Membrane was stained by Ponceau S (0.1% w/v Ponceau S in 5% acetic acid (v/v), afterwards incubated in 3% BSA in TBST (0.1% Tween-20 in Tris-buffered saline) overnight. On the next day, the blot was incubated with 0.3 μg ml^−1^ streptavidin-HRP (Thermo) in TBS-T for 1 hour at room temperature and further developed using the Pierce™ ECL Western Blotting-Substrate (Thermo).

### Streptavidin pulldown for mass spectrometry (MS) sample preparation

Around 1.5 million cells for each replicate were induced with DOX for 24 hours. Controls for uninduced cells and puromycin treatment were seeded at the same time. Next day, cells having 85% confluency were biotinylated and further lysed in the RIPA buffer as described above. For puromycin treatment, cells were treated with 200 μM puromycin and 0.5 mM BP for 30 min at 37°C^16,27^. APEX2 labeling was induced by H_2_O_2_ treatment for one minute as described above. 50 μL streptavidin-coated magnetic beads (Pierce) were used for each replicate and washed three times with RIPA buffer with end-to-end rotation. Pre-washed beads were incubated with 120 μg of lysate for each replicate for 2 hours at room temperature under constant end-to-end rotation. For capturing of biotinylated proteins, we followed a published protocol^34^. After capturing, beads were washed twice with 1 ml of RIPA lysis buffer for 2 min, once with 1 M KCl for 2 min, once with 0.1 M Na_2_CO_3_ for around 10 seconds, once with 2M urea in 10 mM Tris-HCl (pH 8.0) for around 10 seconds, and again twice with 1 ml RIPA lysis buffer for 2 minutes at room temperature. The beads were subsequently transferred to fresh tubes and washed briefly first with RIPA buffer, once with 1ml wash buffer (75 mM NaCl in 50mM Tris-HCL pH 7.5) and then processed for LC-MS/MS analysis as follows.

### Sample preparation for MS

On bead captured interaction partners of TOMM20, TOMM70, or otherwise tagged by NES-, and Mito-APEX2 markers were resuspended in 30 µl of denaturation buffer (6 M urea, 2 M thiourea, 10 mM Tris, pH 8). Disulfide bonds were reduced with 10 mM of DTT for 1h and further alkylated with 55 mM of iodoacetamide for 1h in the dark. Proteins were predigested with 2 µl of LysC (Lysyl endopeptidase; Wako Chemicals) for 3h. For overnight digestion with 2 µl of trypsin (Promega Corporation) samples were diluted fourfold with 50 mM ammonium bicarbonate. On the next day, beads were pelleted, the supernatant was transferred to a new tube and the digestion was stopped by adding 1% of TFA the next day. Contaminants were removed by the PreOmics®’ Phoenix kit prior their submission to the MS.

### LC-MS measurement and data analysis

All proximity labeled samples were analyzed on a Q Exactive HF mass spectrometer (Thermo Fisher Scientific) online-coupled to an Easy-nLC 1200 UHPLC (Thermo Fisher Scientific). Prior to MS-based analysis peptides were separated on an in-house packed (ReproSil-Pur C18-AQ 1.9 μm resin [Dr Maisch GmbH Ltd]) 20 cm analytical column (75 μm ID PicoTip fused silica emitter [New Objective]). For gradient elution of the peptides solvent A (0.1% FA) and solvent B (0.1% FA in 80% ACN) were used across a 60 min gradient with a flow rate of 200 nl/min at 40 °C. The mass spectrometer was operated in positive ion and in data-dependent acquisition mode. MS1 spectra were acquired over a scan range from 300 to 1650 m/z at resolution 60k. The top 7 most abundant peptides were selected for isolation within an isolation window of 1.4 m/z, HCD fragmentation with nce set to 27 and a maximum injection time of 110 ms/ AGC target 1e5. MS2 spectra were acquired at resolution 60k. Peptides were excluded from re-analyzing for 30 s.

Generated raw files were further processed with the MaxQuant software suite^56^, version 2.2.0.0. All parameters were kept as default with trypsin specific digestion mode selected and two missed cleavages allowed. As quantification method, label-free quantification was selected with minimal ratio count of 1. Cystein carbamidomethylation was selected as fixed as well as methionine oxidation and protein N-terminal acetylation as variable modifications. Match between runs was allowed across all raw files. All spectra were searched against the UniProt Homo sapiens database (downloaded on December 16, 2022, 103 830 entries) and commonly observed contaminants.

For downstream data analysis only protein groups were considered that had not been identified as contaminant or by site. Data were analyzed using Perseus software ^57^, version 1.6.7.0 with annotation for cellular compartment and biological function based on gene ontology and MitoCarta3.0^35^.Within biological replicates only proteins quantified in two out of three replicates or if more replicates were performed 75% were used further. To decrease the number of missing values these were replaced from normal distribution with standard settings. To identify significantly higher or lower abundant proteins, a student’s two-sample t-test was performed. Candidates were classified as significant (p-value: 5%, Difference: 3) and stringent significant (p-value: 1%, Difference: 4). Additional data visualization was performed within the R environment. The mass spectrometry proteomics data have been deposited to the ProteomeXchange Consortium via the PRIDE^58^ partner repository with the dataset identifier PXD057097 and 10.6019/PXD057097”.

## Supporting information

Suppl Figures and link to Suppl Tables

## Acknowledgement

We thank Silke Wahl (Proteome Center Tübingen) for sample processing and Bianca Lemke (Proteome Center Tübingen) for technical support in data processing. We also thank Julien Béthune (HAW Hamburg) for providing plasmids and cell line. We thank Birgit Singer-Krüger (Interfaculty Institute of Biochemistry, Tübingen) for providing APEX2 antibody. We also thank Gabriel Dodt (Interfaculty Institute of Biochemistry, Tübingen) for lab equipment and reagents. We also thank to Nisha Mohd Rafiq (Interfaculty Institute of Biochemistry, Tübingen) for sharing lab equipment, lab reagents, and technical support for microscopy analysis, and Moritz Lehners (Interfaculty Institute of Biochemistry, Tübingen) for technical support to handle microscope. SA and KIZ were supported by funding via the GRK2346 “MOMbrane” of the Deutsche Forschungsgemeinschaft.

## Author Contributions

SA, KIZ, RPJ wrote the paper with the support of BM, SA designed the project and performed all experiments. KIZ and BM were responsible for MS sample processing and data analysis. Figures were made by SA and KIZ.

