## Supplementary material for "Proximity labeling reveals differential interaction partners of the human mitochondrial import receptor proteins TOMM20 and TOMM70": Suppl Figures and link to Suppl Tables

**List of Supplementary figures and tables**


| | |
|---|---|
| Supplementary Figure 1 | Expression of cytoplasmically (NES-APEX2) and mitochondrial matrix localized (mito-APEX2) fusion proteins |
| Supplementary Figure 2 | Compartment specific local biotinylation mediated by NES-APEX2 and mito-APEX2 |
| Supplementary Figure 3 | Quantitative analysis of TOMM20- and TOMM70-APEX2 interactomes |
| Supplementary Figure 4 | Representative confocal images of immunostaining showing colocalization of SYNJ2BP with TOMM20 |
| Supplementary Figure 5 | Comparisons of proteins identified in various interactomes and effect of puromycin on TOMM20- and TOMM70-APEX2 interactomes |
| Supplementary Table 1 | Proteins identified in TOMM20-APEX2 vs -DOX interactome |
| Supplementary Table 2 | Proteins identified in TOMM70-APEX2 vs -DOX interactome |
| Supplementary Table 3 | Proteins identified in TOMM20-APEX2 vs NES-APEX2 interactome |
| Supplementary Table 4 | Proteins identified in TOMM70-APEX2 vs NES-APEX2 interactome |
| Supplementary Table 5 | Proteins identified in TOMM20-APEX2 vs Mito-APEX2 interactome |
| Supplementary Table 6 | Proteins identified in TOMM70-APEX2 vs Mito-APEX2 interactome |
| Supplementary Table 7 | Proteins identified in TOMM20-APEX2 vs TOMM70-APEX2 interactome |
| Supplementary Table 8 | Significantly enriched proteins of TOMM20-APEX2 identified after comparing TOMM20-APEX2 vs TOMM70-APEX2 interactome |
| Supplementary Table 9 | Overlapping of TOMM20-APEX2 candidates among different interactomes |

| | |
|---|---|
| Supplementary Table 10 | Proteins identified in TOMM20-APEX2 vs TOMM20-APEX2 (+puro) interactome |
| Supplementary Table 11 | Proteins identified in TOMM70-APEX2 vs TOMM70-APEX2 (+puro) interactome |
| Supplementary Table 12 | Significantly enriched proteins of TOMM20-APEX2 (+puro) identified by comparing TOMM20-APEX2 (+puro) vs TOMM70-APEX2 (+puro) interactome |
| Supplementary Table 13 | Proteins identified in TOMM20-APEX2 (+puro) vs TOMM70-APEX2 (+puro) interactome |
| Supplementary Table 14 | List of plasmids generated in this study |
| Supplementary Table 15 | List of antibodies used in this study |

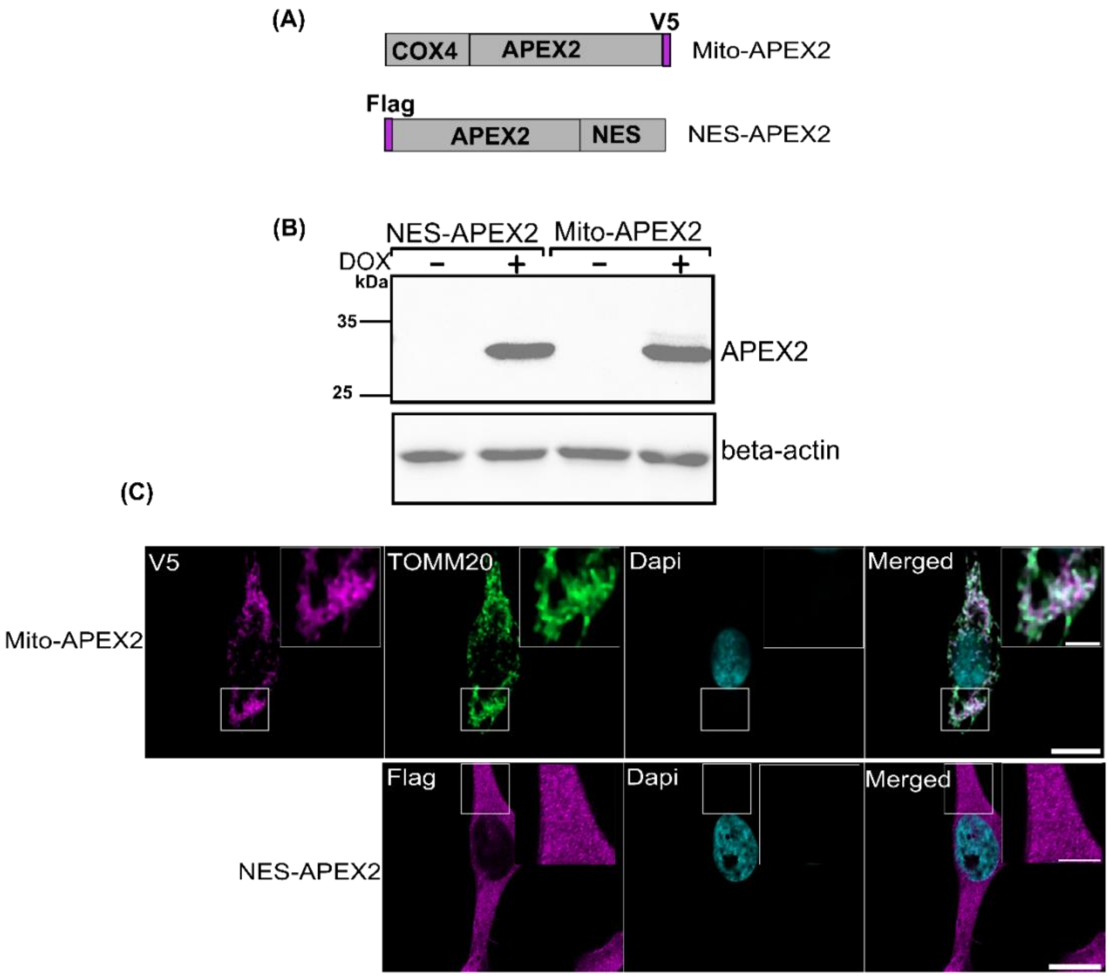


**Supplementary Figure 1. Expression of cytoplasmically (NES-APEX2) and mitochondrial matrix localized (mito-APEX2) fusion proteins**

**(A)** Domain structures of fusion proteins stably expressing APEX2 in various cellular compartments. Mito-V5-APEX2 harnesses a 1-24 amino-acid sequence from Mitochondrial matrix resident 'COX4' protein to localize APEX2 in the mitochondrial matrix. FLAG-APEX2-NES utilizes a nuclear export signal (NES) to target APEX2 in the cytoplasm.

**(B)** Western blot analysis of whole cell lysate of cells stably expressing Mito-APEX2, and NES-APEX2 constructs. Cells are either not or induced with DOX for 24 hours prior to lysis. APEX2 containing fusion proteins are analyzed with APEX2 antibody. beta-actin is used as loading control.

**(C)** Confocal fluorescence imaging confirming the cellular localization of stably expressing Mito-APEX2 and NES-APEX2 fusion proteins. Cells are induced with DOX for 24 hours, immunolabeled with antibodies directed against V5 to detect Mito-APEX2 proteins (Magenta), or flag to visualize APEX2-NES fusion protein (Magenta). (Scale bar: 10 μm; incent: 5 μm).

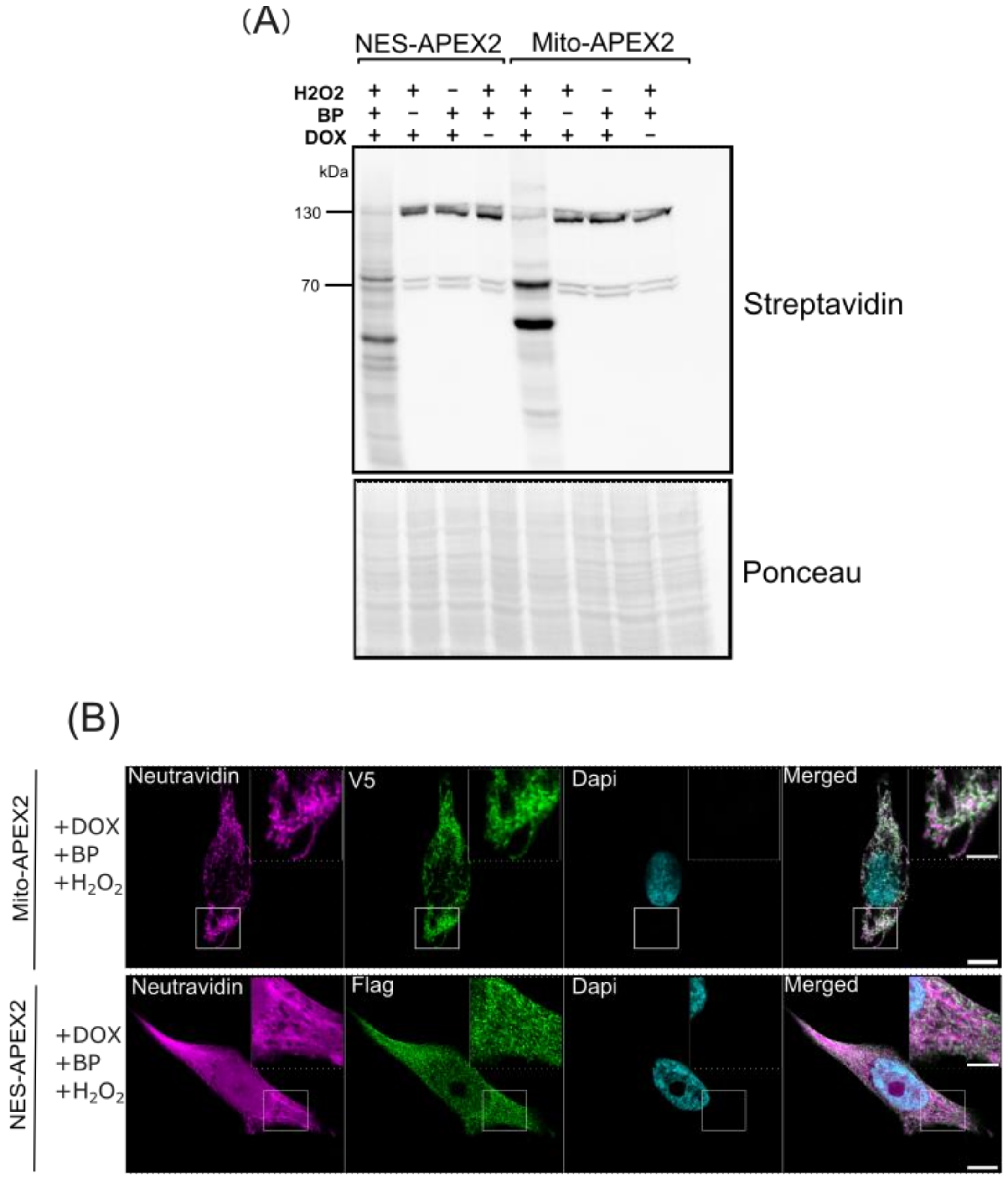


**Supplementary Figure 2. Compartment specific local biotinylation mediated by NES-APEX2 and mito-APEX2**

**(A)** Western blot analysis of cell lysate to confirm the APEX2-mediated biotinylation in the cells stably expressing Mito-APEX2 and NES-APEX2 constructs. Biotinylated proteins are probed by Streptavidin-HRP conjugate (upper portion), ponceau staining is shown on the lower portion.

**(B)** Confocal fluorescence imaging of APEX2-mediated biotinylation in Hela cells stably expressing the Mito-APEX2 and NES-APEX2 constructs. Following 24 hours of DOX induction, cells are subjected to live-cell biotinylation with Biotin phenol (BP) and $H_2O_2$ for one minute, then fixed. Cells are subsequently stained with either 'V5' or 'flag' antibodies to confirm the expression of the indicated APEX2 fusion proteins (appearing in green), while 'Neutravidin' is utilized to stain the biotinylated species (appearing in magenta). Nuclei are stained with DAPI (cyan). (Scale bar: 10 μm; incent: 5 μm).

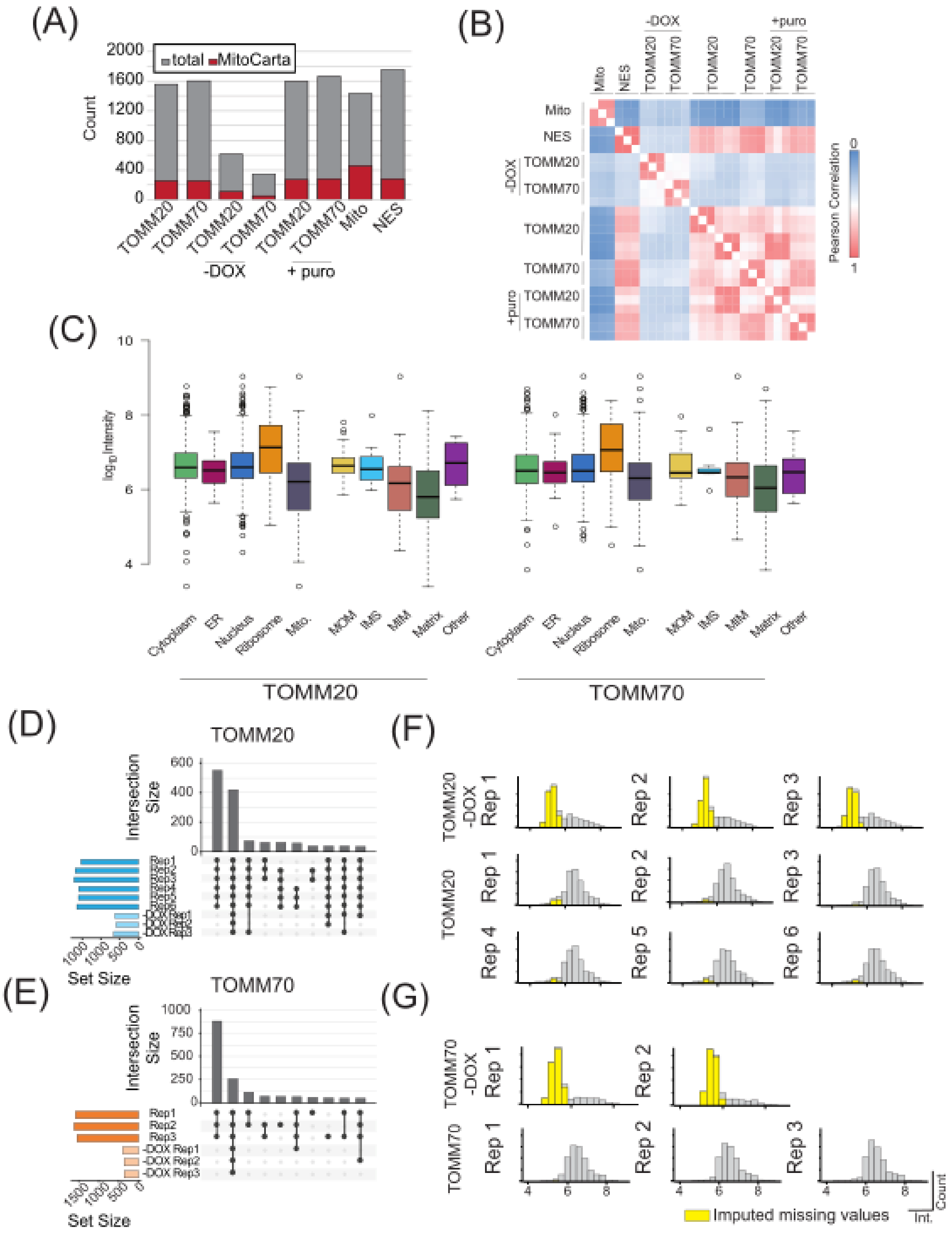


**Supplementary Figure 3. Quantitative analysis of TOMM20- and TOMM70-APEX2 interactomes**

**(A)**Total identification of proteins (grey) and proteins annotated for mitochondrial localization (red) based on MitoCarta3.0. Count of proteins based on quantification in minimum 3 out of 6 replicates for TOMM20-APEX and 2 out of 3 replicates for remaining samples. (**B)** Correlation between replicates based on Pearson correlation. (**C)** Box plots showing the log-transformed intensity of organelle-annotated proteins identified in TOMM20-APEX2 (left) and TOMM70-APEX2 (right). (**D)** Upset plot of overlapping proteins identified between replicates of TOMM20-APEX2 and controls (-DOX TOMM20-APEX2). (**E)** Upset plot of overlapping proteins identified between replicates of TOMM70-APEX2 and controls (-DOX TOMM70-APEX2). Imputation of missing values (yellow bars) showing mostly unidentified low abundant proteins replaced from normal distribution for -DOX TOMM20-APEX **(F)** and -DOX TOMM70-APEX2 **(G).**

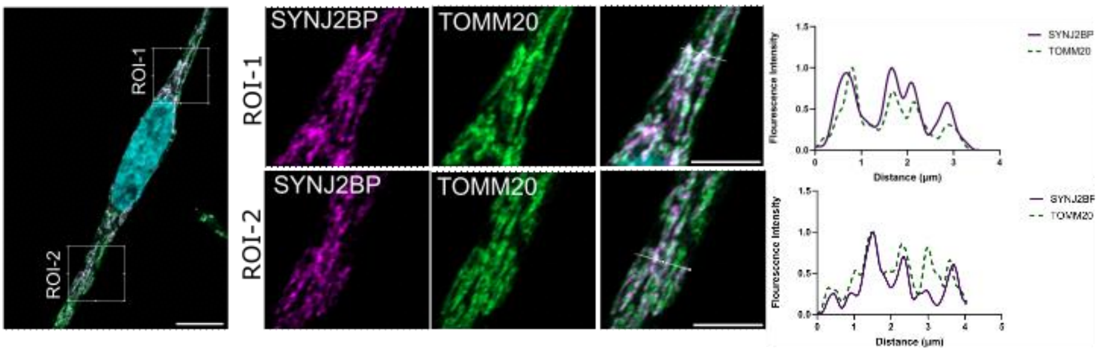


**Supplementary Figure 4. Representative confocal images of immunostaining showing colocalization of SYNJ2BP with TOMM20**

Super resolution images showing co-localization of SYNJ2BP to TOMM20. Hela cells are fixed and immunolabeled with antibodies directed against TOMM20 (Green) and SYNJ2BP (Magenta). Each representative confocal image represents a merged image of green and magenta channels, and two ROI are selected in the cell. Nuclei are stained with DAPI (cyan) (Scale bar: 10 µm). Each ROI is a zoomed-in portion of the cell showing in the middle (Scale bars, 5 µm). Scan intensity profiles represent the overlapping fluorescence signals of SYNJ2BP and TOMM20.

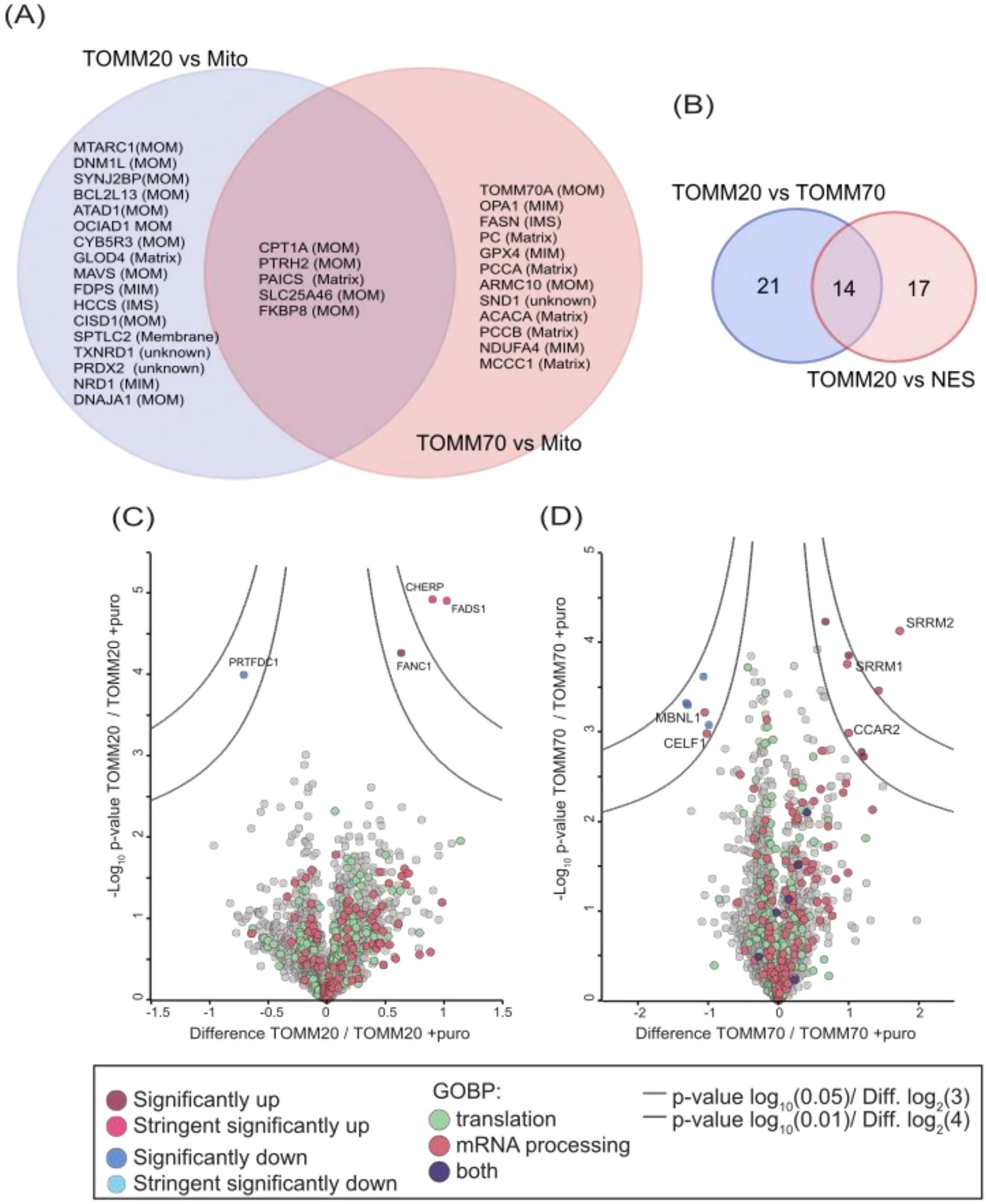


**Supplementary Figure 5. Comparisons of proteins identified in various interactomes and effect of puromycin on TOMM20- and TOMM70-APEX2 interactomes**

**(A)** Venn diagram showing overlapping between the proteins enriched in TOMM20-APEX2 vs Mito and TOMM70-APEX2 vs Mito. (**B)** Venn diagram showing TOMM20 overlapping candidates by comparing the enriched candidates of TOMM20 in TOMM20 vs TOMM70 interactomes and TOMM20 vs NES interactome. (**C)** Volcano plots for TOMM20-APEX2 interactome against TOMM20-APEX2 interactome (+puro). (**D)** Volcano plots for TOMM70-APEX2 interactome against TOMM70-APEX2 interactome (+puro). Highlighted are the proteins annotated (based on GOBP) for translation and mRNA processing.

Supplementary tables 1 – 13 (Excel files) and 14 + 15 can be accessed online via the following link: https://drive.google.com/drive/folders/1DRhP1r0_h7EoKaV4uxoSXHOgLpir7tlZ?usp=sharing
